# A network perspective on the evolution of hybrid incompatibilities

**DOI:** 10.1101/2025.07.09.663985

**Authors:** Evgeny Brud, Rafael F. Guerrero

## Abstract

Theory predicts that hybrid incompatibilities accumulate faster than linearly with genetic divergence, a phenomenon known as the snowball effect. While this prediction is mathematically robust under simplifying assumptions, accumulating evidence suggests that the structure of gene interaction networks can alter both the rate and organization of incompatibility evolution. Here, we extend classic DMI models with a network approach, equating the assumptions of the Orr model with a complete graph of gene interactions. We simulate the evolution of hybrid incompatibilities under different gene interaction networks and evaluate the effects of network density, topology, and substitution model. We find that network density strongly governs the rate of DMI accumulation, particularly under models permitting multiple substitutions per locus, while network topology shapes the agglomeration of incompatibilities into large, connected clusters. Substitution rate heterogeneity, especially when anti-correlated with node degree, further suppresses both accumulation and clustering. These results highlight that while the snowball effect remains qualitatively valid, the structure and evolution of the incompatibility network exhibit nontrivial departures from previous expectations, with implications for observable quantities in empirical systems. Our findings underscore the importance of incorporating genomic architecture and network constraints into models of speciation.

## Introduction

A central question in speciation genetics is how reproductive isolation arises from the accumulation of genetic differences between diverging populations. Postzygotic isolation is often a product of independent substitutions in divergent populations that, when combined in hybrids, result in deleterious epistatic interactions (i.e. Dobzhansky– Muller incompatibilities, or DMIs; Dobzhansky, 1936; Muller 1942). Orr (1995) formalized this idea and showed that, under simple assumptions, the number of pairwise DMIs grows at least quadratically with divergence, thereby producing a “snowball effect.” In this model, two lineages accumulate substitutions at *K* loci, and each new allele has a small probability *P* of interacting negatively with any substituted allele in the other genome. As divergence proceeds, the number of potential incompatibilities increases combinatorially, causing the probability of reproductive isolation to accelerate with time.

Subsequent work has extended Orr’s framework, finding that, while the overall superlinear expectation is robust to several key assumptions, genome structure determines the pattern of DMI accumulation. For instance, allowing a modular genome or relaxing assumptions of symmetric substitution rates does not have a qualitative effect on the expected snowball pattern (Turelli and Orr 2000; Orr and Turelli 2001). More recent approaches modeling divergence in RNA-folding and protein-protein interaction networks have relaxed the assumptions of infinite genome size, low probability of incompatibility, and the omnigenic model (by restricting interactions to connected loci). In RNA folding landscapes, the rapid accumulation of simple pairwise DMIs gives way to the formation of higher-order incompatibilities, a process termed *complexification*, which slows the apparent rate of accumulation (Kalirad and Azevedo 2017). In protein-protein interaction networks, density has been shown to be a key determinant of the rate of accumulation (Livingstone et al. 2012). These findings collectively suggest that the underlying interaction network plays a central role in shaping both the tempo and form of DMI accumulation and must be accounted for in models of speciation.

The potential for complex hybrid incompatibilities observed in natural systems reinforces the need for more realistic models (reviewed in Satokangas et al 2020; Ayala-Lopez and Bank 2024). In *Drosophila* hybrids, inviability is linked to multi-locus interactions that often involve the sex chromosomes (Cabot et al. 1994; Palopoli & Wu 1994; Tang & Presgraves 2009; Phadnis et al. 2015). In *Solanum*, double-introgression studies have shown widespread antagonistic epistasis between loci involved in pollen sterility, implying that the barrier hinges on the joint presence of multiple introgressions and suggesting that highly interconnected incompatibility networks underlie isolation (Moyle & Nakazato 2009; Sherman et al 2014; Guerrero et al. 2017). In *Xiphophorus* fishes, a recent large-scale genetic dissection of hybrid sterility found that it arises from non-additive interactions among three or more loci dispersed across the genome (Moran et al. 2024). Further evidence of complex DMIs has been found, albeit indirectly, in *Nasonia* wasps (Koevoets et al 2012), *Formica* ants (Heidbreder et al 2025), and *Saccharomyces* (Kao et al 2010). These observed patterns may help explain the mixed empirical support for a simple snowball expectation (Matute et al. 2010; Moyle and Nakazato 2010; Kondrashov et al. 2002; Wang et al. 2015) and underscore the importance of modeling DMIs within the context of explicit genomic structures (Ayala-Lopez and Bank 2024).

To address how genomic architecture shapes the evolution of reproductive isolation, we extend classical DMI models by embedding them within gene interaction networks (Fig. 1). We represent genomes as undirected graphs, where nodes are loci and edges represent potential interactions. Our aim is to quantify how network density (the relative abundance of edges) and topology (the structure of edge connections) influence the formation of the incompatibility network. We simulate DMI evolution for various network topologies: (i) scale-free (Barabási and Albert 1999), (ii) small-world (Watts and Strogatz 1998), (iii) complete graphs and (iv) a digenic interaction network from yeast (Costanzo et al. 2016). We track the number of nodes and edges on the incompatibility network over time (the unit of time being a substitution event). We also examine how different substitution regimes (i.e. limiting each locus to a single substitution or allowing multiple hits) affect DMI formation. For scale-free and yeast networks, we further incorporate differential rates of evolution among loci, in which some loci deviate markedly in their number of interactions as compared to the typical locus and consequently may exhibit wide variation in substitution rates (e.g. constraints on allelic replacement at hub genes owing to their greater likelihood of deleterious pleiotropy).

**Figure 1.**
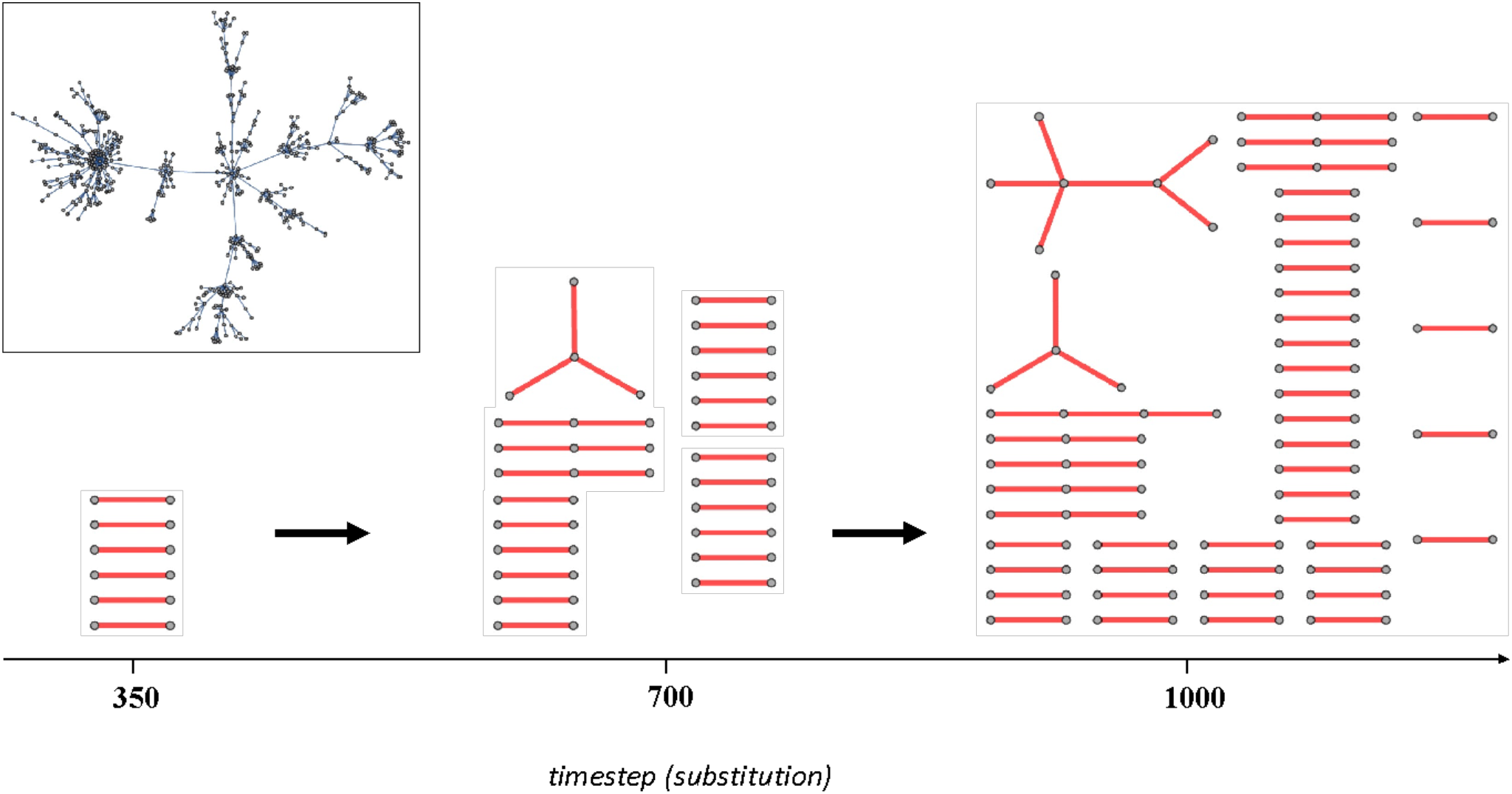
Illustration of pairwise incompatibility evolution. Assuming a network of gene interactions (inset), populations in allopatry build up a network of pairwise incompatibilities over time. Typically, early stages of speciation involve the accumulation of incompatible pairs. Over time, DMI pairs develop connections to additional loci and/or clusters, resulting in connected components of greater size and complexity than isolated edges.

A core focus of our work is the concept of agglomeration, quantified by tracking the size distribution of connected DMI components. In addition to increasing in number, incompatibilities may link to existing DMIs through shared interacting loci, forming larger and more interconnected clusters. This process, analogous to complexification (Kalirad and Azevedo 2017), creates higher-order structures in the incompatibility network that differ qualitatively from isolated pairs and may alter predictions about the rate and genetic architecture of speciation. We demonstrate that the substitution model, the probability of digenic incompatibility, and features of gene-interaction networks, such as density and topology, interact to determine the rates of DMI accumulation and agglomeration. In particular, sparse interactions (where vertices and edges of the underlying network are of the same magnitude) constrain DMI accumulation, while both sparseness and small-world topology constrain DMI agglomeration. Variation in per-locus substitution rates, when negatively correlated to the number of interactions, adds an additional and potentially severe constraint on agglomeration and thereby promotes the accumulation of small DMI complexes.

## Methods

We describe the genome as an undirected network of interactions (edges) between loci (nodes). The loci corresponding to the fixed differences separating a pair of populations, along with their full set of associated interactions, constitutes an induced subgraph of the genome-wide interaction network. A subset of the edges from this induced subgraph constitutes a pairwise incompatibility network that contains only edges and nodes involved in incompatibilities. Orr’s (1995) combinatorial treatment is readily discerned as equivalent to assuming a complete network of gene interactions with equal probability of incompatibility (*P*). Small values of *P* are sensible in this context of omnigenic interaction, given that a maximal density of edges is present; as Orr (1995) estimated, a value of *P* = 10^−6^ was roughly consistent with contemporary evidence from *Drosophila* (i.e. of sufficient magnitude to yield a single DMI given the divergence of a known species pair). However, *P* and the number of substitutions are, in general, not a sufficient set of parameters with regard to the numerical evolution of pairwise DMIs. This deficiency partly follows from (1) the fact that only a subset of hybrid genotypic combinations involve loci that interact in the parent species; a *P* of zero should be assumed in the absence of interaction (Livingstone et al. 2012), and (2) assumptions regarding the correlation between genome-wide substitution rates and the degree distribution are certain to affect evolutionary outcomes (Olofsson et al. 2016). Consequently, far higher values of *P* are explored here when investigating networks with sparse connections between loci (see Kondrashov et al. 2002, Welch 2004 for data and theory supporting a high *P*).

### Gene interaction networks

Gene interaction networks with 1,000 nodes were generated using the Watts-Strogatz Graph Distribution (small-world, rewiring p = 0.1, k-parameter ∈ {1,5}), Barabási-Albert Graph Distribution (scale-free, k-parameter ∈ {1,5}), and Complete Graph functions in Mathematica (v13.0, Wolfram Research Inc. 2021), with 250 replicates (excepting complete graph). The yeast gene interaction network was obtained from Costanzo et al. (2016) and filtered according to the authors’ recommendations for a stringent setting (significance *p*-value < 0.05 and interaction-score > 0.16 or < -0.12); the resulting network (5841 nodes, 421628 edges) was further pared by only including loci with available dN/dS values from Peter et al. (2018), resulting in a network with 5502 nodes and 367908 edges.

### Incompatibility accumulation regimes

For our simulations, we assumed three regimes of incompatibility build-up, as determined by the product of *P* and the density of digenic interactions (**low, medium, high**; Table 1). Gene interaction networks with different graph densities were assigned “density-adjusted” *P* values in order to fix the overall sensitivity to incompatibility origination associated with a given regime (Table 1). For the multiple-hits scenarios, locus pairs may build up multiple edges and so equality of DMI-edge totals (counting each multi-edge) over K substitutions defines two regimes as having equivalent incompatibility regimes (equality of single-edges is a sufficient criterion under single-hits-only scenarios); simple-graph edge totals are displayed in all edge-count trajectory figures. In particular, we assumed *P* values of 0.01%, 0.06%, and 0.16% for the low, medium, high regimes for the complete network, respectively; we then adjusted the remaining *P* values (for 0.2%-density, 1%-density, and Yeast networks) based on the ratios of graph densities.

**Table 1.** Probabilities of incompatibility per digenic interaction assumed in simulations.

| Incompatibility regime | Complete<br>(1,000 nodes \| 499,500 edges) | 0.2%-density* | 1%-density** | Yeast network<br>(5,502 nodes \| 367,908 edges) |
| --- | --- | --- | --- | --- |
| **Low** | 0.0001 | 0.05 | 0.01 | 0.0041 |
| **Med** | 0.0006 | 0.30 | 0.06 | 0.0246 |
| **High** | 0.0016 | 0.80 | 0.16 | 0.0656 |

\* scale-free: 999 edges; small-world: 1,000 edges; 1,000 nodes, both. Densities are approx. equal.
\*\* scale-free: 4,985 edges; small-world: 5,000 edges; 1,000 nodes, both. Densities are approx. equal.

### Computer simulations

Substitution histories were created (250 replicates) by sampling nodes with replacement (multiple substitutions) for 1500 timesteps or without replacement (single-hit substitutions) for 1000 timesteps. Substitution histories were re-used for the 0.2%, 1%, and Complete network scenarios. Yeast substitution histories were similarly created with nodes corresponding to systematic gene names. Differential evolutionary rates among loci were incorporated by including a vector of weights with the Random Choice function in Mathematica. For 0.2%, 1% and Complete network scenarios, the vector of weights equaled the degree distribution raised to an exponent (−*α*) where *α* ∈ {0, 0.5, 1, 2}. For differential selection *random with respect to connectivity*, the vector of weights was randomized by order. Differential evolutionary rates in yeast were incorporated by setting the vector of weights equal to the vector of corresponding dN/dS values (Peter et al. 2018) and compared to simulations where the ordering of weights was randomized.

For each substitution history, a network of pairwise incompatibilities was constructed iteratively. At each timestep *k*, the edges incident to N_*k*_on the induced subgraph containing N_1_, …, N_*k*_were Bernoulli sampled with probability *P*, where N_*k*_gives the node corresponding to the substitution at timestep *k*.

### Data availability

All interaction networks, substitution histories, incompatibility networks, and statistical analyses are available as “.wl” files (Wolfram Mathematica).

## Results

### Patterns of DMI accumulation

With multiple substitutions permitted per locus (assuming equal substitution probabilities), network density has a sizeable effect on the accumulation of edges in the medium and high accumulation regimes (Fig. 2). This is because loci subjected to multiple substitutions not only contribute to incompatibility *origination*, but also to incompatibility *turnover*. That is, some locus-pairs repeatedly evolve an incompatible interaction at two or more time-steps; these turnover events do not add novel edges or nodes to the pairwise incompatibility network. Low-density gene interaction networks (small-world-0.2%, scale-free-0.2%; each with ∼1000 edges) devote a larger fraction of Bernoulli trials to turnovers, which reduces their rates of edge accumulation compared to more interaction-rich networks. The slower accumulation is more pronounced for DMI edges than for DMI loci (Fig. 2). Comparison across the denser networks (small-world-1% vs. scale-free-1% vs. complete) reveals that (i) topology plays only a modest role and (ii) the effect of density differences is considerably reduced when comparing ∼5000-edge networks with complete networks (∼500,000 edges) (Fig. 2). Overall, the incompatibility regime and the lower end of density variation are the dominant factors determining the increase of DMI interactions.

**Figure 2.**
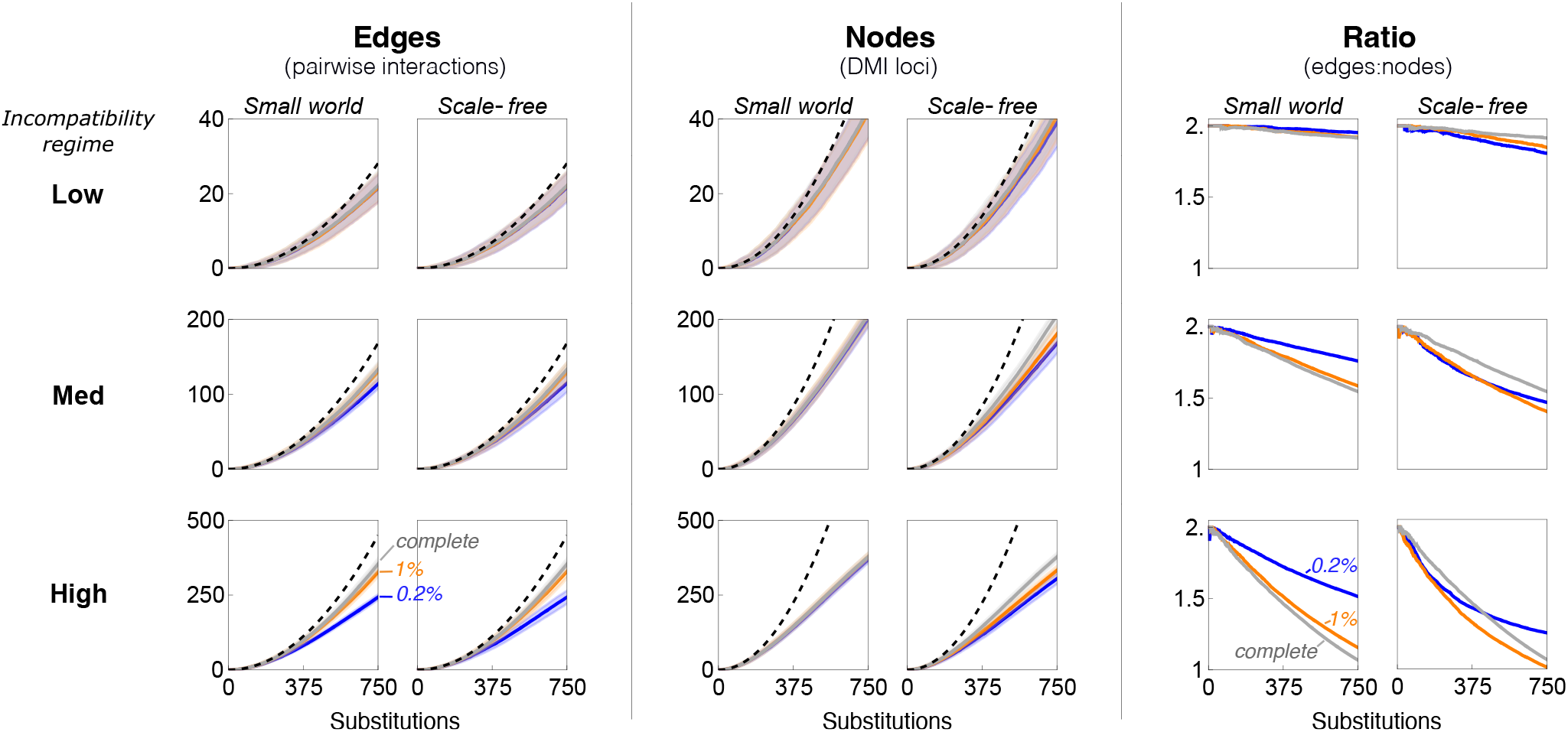
Incompatibility accumulation under a multiple-hits, equal rates substitution model. Counts of edges (left), nodes (middle), and the ratio of those two values (right) on the pairwise incompatibility network under low, medium, and high DMI accumulation regimes. Evolution occurred assuming small-world and scale-free gene interaction networks with 1000 loci. Density of the underlying interaction network is given by color (blue: 0.2% density; orange: 1% density; gray: complete network). Complete graph trajectories are duplicated in corresponding small-world and scale-free panels for edges and nodes. Solid lines are mean values. When present, bands show ± 1 mean deviation. Dashed lines plot Orr’s (1995) binomial expectation for DMI incompatibilities (edges), and for DMI loci based on a 2:1 ratio of loci to edges.

The accumulation results are also reflected in the evolving ratios of DMI loci to DMI interactions (Fig. 2). Initial adherence to a 2:1 ratio declines linearly under the low DMI rate for all cases; the decline is faster under medium to high DMI rates and often nonlinear. The evolution of ratios below 1 is a common outcome under a high incompatibility regime.

Assuming a single-hit substitution model, the rate of incompatibility increase is again determined strongly by the incompatibility regime, but here the influences of network density and topology are both slight (Fig. S1). The singlehit model maintains a consistent rate of snowballing (i.e. a constant acceleration of DMIs per unit time, ignoring early noise in the first ∼200 timesteps; Fig. S2), while DMI accumulation in the multiple-hits model decelerates over time owing to the turnover effect.

### Patterns of DMI agglomeration

As mentioned, the aggregation of edges into connected components of increasing size occurs alongside the numerical increase of DMIs. One measure of this process is given by the number of connected components present on the incompatibility network (Figs. 3, S3a-f). Notably, the trajectory of component counts is seen to be *non-monotonic*, so that while a snowballing increase is initially observed, a precipitous decline appears under medium and high DMI accumulation rates (Figs. 3, S3c-f). A look at the size distribution reveals that the cause of this reversal owes to the agglomeration of connected components into fewer and fewer clusters of increasing size, so that over the long run, a single giant component manifests on the incompatibility network (Erdos and Renyi 1960), alongside a decreasing fraction of pairs and small DMI complexes that make up the remainder of DMIs (Figs. S4-5).

**Figure 3.**
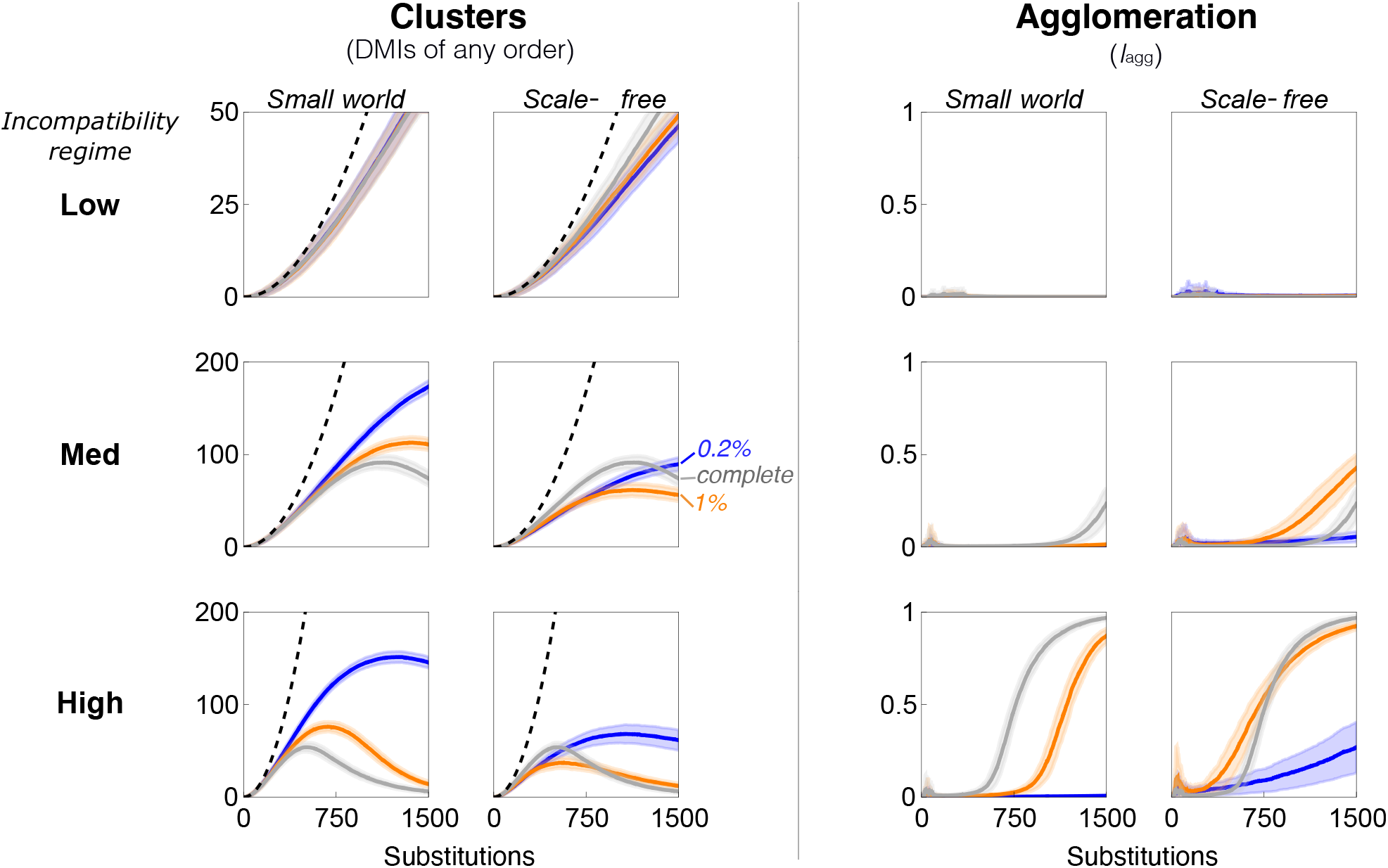
Incompatibility agglomeration under a multiple-hits, equal rates model. Density of the underlying interaction network is given by color (0.2% density – **blue**; 1% density – **orange**; Complete – **gray**). Complete graph trajectories are duplicated in corresponding small-world and scale-free panels. Left: the number of connected components on the pairwise incompatibility graph (means ± 1 mean deviation). Dashed line corresponds to Orr’s (1995) binomial expectation for DMI incompatibilities. Right: *I*_agg_(t), index of agglomeration (means ± 1 mean deviation).

Consequently, we desire a statistic that distinguishes between a network consisting mostly of isolated DMI pairs and small complexes on the one hand and a DMI network consisting largely of a single massive cluster on the other. We define the following *index of agglomeration*, where *c*_*t*_is the number of connected components at time *t* and *L*_*C*(i,*t*)_ is the size of the i^th^ connected component at *t*:

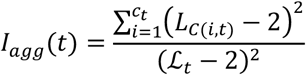

where ℒ_*t*_ is the number of DMI loci at time *t*. The numerator is the squared Euclidean distance between the point 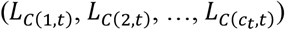 and a reference (2, 2, …, 2), where the 2’s are repeated *c*_*t*_ times. The index is therefore a measure of divergence between an observed DMI network and a reference network comprised of strictly pairwise DMIs (where all vertex degrees are equal to one) containing exactly as many connected components as the observed network. Note that the motivation for squaring the distance is to emphasize large deviations. *I*_*agg*_(*t*) is a normalized measure, taking values between 0 and 1.

We compare agglomeration trajectories and see that network topology and density both drastically affect the clustering process (Figs. 3, 4). In particular, lower network density and a small-world topology are each generally associated with much slower rates of increase in *I*_*agg*_ (*t*) (Fig. 3). Under our “medium” incompatibility regime (Table 1), scale-free networks cluster more rapidly than complete networks, while under a “high”-incompatibility regime, their course of agglomeration is similar.

**Figure 4.**
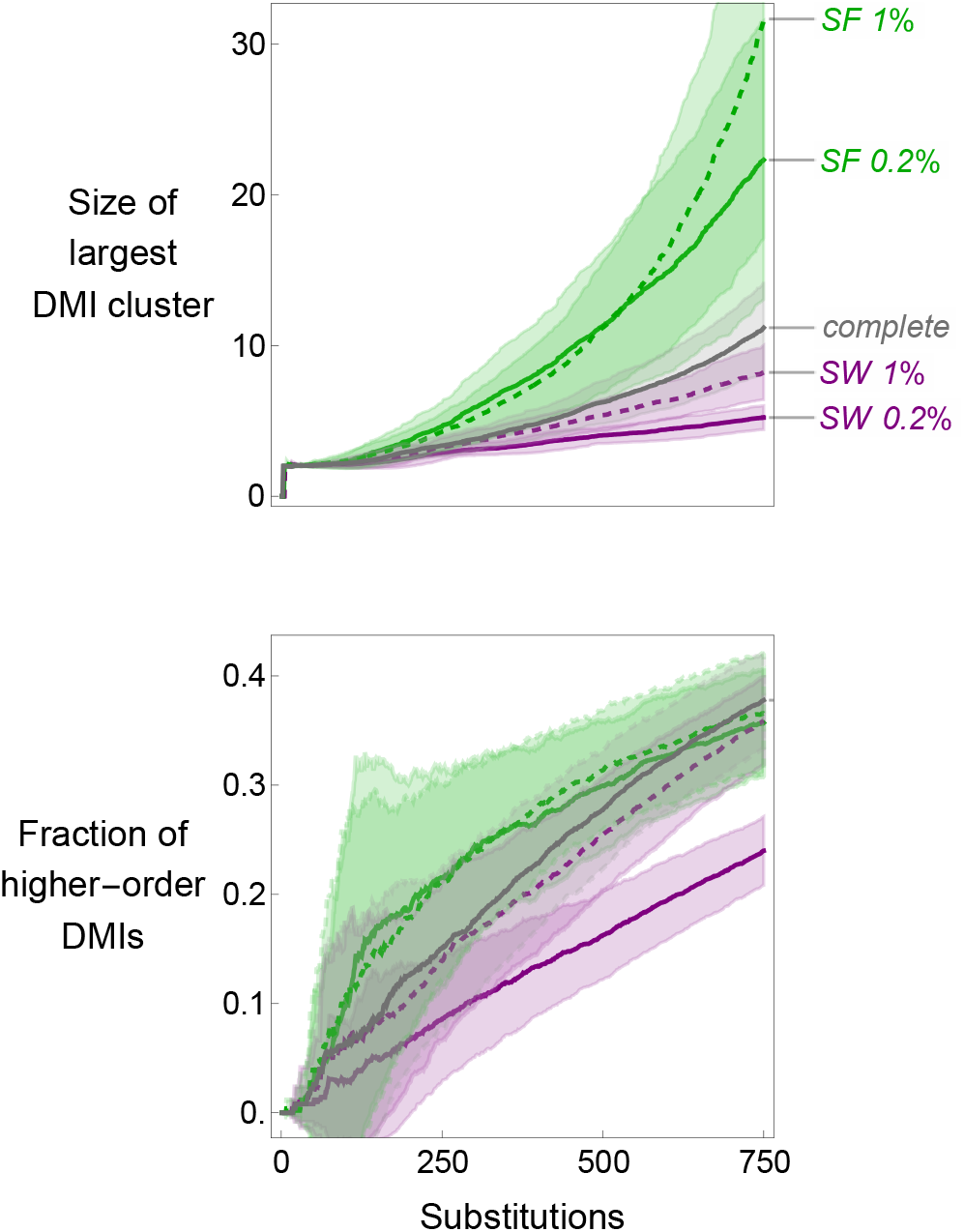
Evolution of incompatibility clusters. The size of the largest observed DMI cluster (top panel) is heavily dependent on the underlying network topology (SF=scale-free, SW=small-world; shown at two densities). The fraction of all DMI clusters that are composed of three or more loci (‘higher-order; bottom panel) is affected by the interaction of density and topology. Simulations run under a *medium* DMI accumulation regime, assuming multiple-hits, equal substitution rates model. Plots for *low* and *high* regimes are qualitatively similar (in the Supplement). Bold lines are means banded by ± 1 mean deviation, conditional on replicates having > 0 incompatibilities.

An additional statistic for which the scenarios display differentiated trajectories is the maximum degree among the nodes on the DMI network. Scale-free networks at either density (0.2% or 1%) experience a steady increase in the degree of the most highly connected DMI locus (an incompatibility hub), while the rate of increase in this quantity increases at a far slower rate (∼2x-10x slower) for small-world and complete networks under all three incompatibility regimes (Fig S6). This is closely reflected in the size evolution of the largest DMI cluster (Fig. 4).

### Differential rates of evolution among loci

Restricting our attention in this section to scale-free topology, we report the influence of substitution rate variation on the pace of DMI evolution. A uniform substitution rate across the genome (*α* = 0) always produces the fastest accumulation of DMI loci (Fig. 5A). Assuming that substitution rates vary, and moreover, may be correlated with the degree distribution of the underlying interaction network, we compared two scenarios: (1) “Mutation by degree” -where more highly connected genes are stipulated to have *lower* substitution rates according to the power law distribution described in Methods, and (2) “Random variance” - where these substitution rates are reused but randomized across nodes. A strong negative correlation between evolutionary rates and connectivity is observed to slow the accumulation of DMI loci by a factor of up to 4-fold (*α* = 2 vs *α* =0, Mutation by degree column). The absence of a strong negative correlation erases this distinction, even in the presence of wide rate variation (compare *α* = 2 between “Random variance” and “Mutation by degree” panels). We conclude from these results that the intensity of rate-variation and the degree-rate correlation interact to determine the rate of the snowball effect. Likewise, this coupling of factors affects the snowball pattern by markedly slowing the process of agglomeration, reflecting both the overall slower pace of DMI inputs into the clustering process as well as the strong negative selection against hub-like nodes from entering the pairwise incompatibility network (Fig. S7).

**Figure 5.**
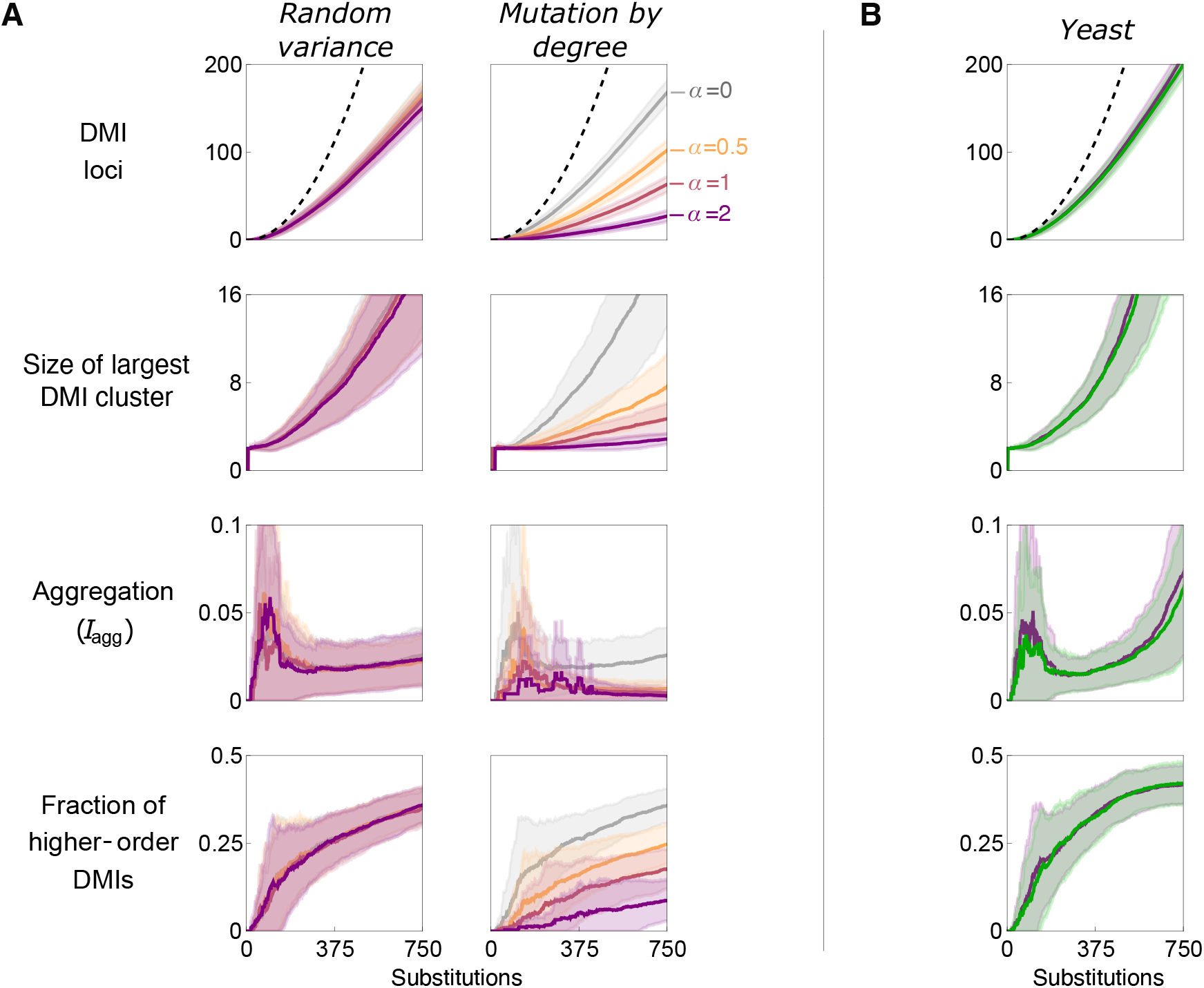
DMI evolution under differential substitution rates. Simulation results (means ± 1 mean deviation) assuming scale-free networks with 0.2% (left) and 1% (right) density are displayed. (A) Trajectories assume values of 0 (equal rates), 0.5 (weak differences in rates), 1 (moderate differences), and 2 (strong differences). Substitution rates are negatively correlated with degrees (“Mutation by degree” column) or are otherwise randomly associated (“Random variance”). Size of largest cluster and fraction of higher-order DMIs are conditional means/mean deviations (for replicates with >0 DMIs). Dashed line for DMI locus panel is based on the 2:1 expectation from the edge-count. (B) Trajectories for DMI evolution assuming a yeast interaction network were simulated under a multiple-hits substitution model; the bands strongly overlap for the two cases depicted: equal substitution rates (purple) and substitution rates proportional to dN/dS values (green).

## Discussion

Our results show that the long-run accumulation and organization of Dobzhansky–Muller incompatibilities (DMIs) are strongly shaped by properties of the underlying gene interaction network. Consistent with previous results (Livingstone et al 2012), network density exerts a dominant influence on the rate of DMI accumulation, particularly for our multiple-hits model. As DMIs agglomerate into larger connected components, network topology plays an increasingly important role modulating the pace and structure of the clustering process. In the absence of strong negative correlations between rates of molecular evolution and the number of gene interactions per locus, scale-free networks give rise to highly connected incompatibility hubs, while small-world and complete graphs show greater uniformity in the connections among DMI loci. Substitution rate heterogeneity can substantially suppress both accumulation and agglomeration (especially when strongly anti-correlated with network connectivity), underscoring the importance of evolutionary constraints imposed by network architecture.

While network effects were mainly observed at later stages of divergence (e.g. *K* > 500 substitutions), it should be noted that the probability of incompatibility (*P*) modulates the time necessary to reach the stage at which the edges of the incompatibility network have accreted into larger and fewer components. For large enough *P* and *K*, the build-up of deleterious hybrid gene-combinations between a pair of populations necessarily exceeds the regime over which one or merely a few DMIs prevent gene flow, as well as the regime over which a simple 2:1 extrapolation of DMI loci to DMI interactions holds approximately true. Moreover, to the extent that the course of fitness decline in hybrids (not modeled in the present article) is protracted rather than single-step or occurring over a small number of substitutions (see Orr and Turelli 2001), the network effects and substitution models will together determine the evolving pattern of fitness interactions as mediated by the hybrid incompatibility network.

These results extend classic expectations from DMI theory. Orr (1995) derived an expectation for the number of DMI interactions, showing that 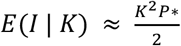, where *I* is the number of DMIs (see Orr and Turelli 2001 for a continuous-time treatment). In the case of omnigenic interaction (complete network), *P*\* is small and uniform; in less-interactive genomes, *P*\* can be interpreted as a *marginal probability,* equal to *P* times the network density (Table 1). In our comparison of substitution models (i.e. loci sampled with or without replacement), the simulation results demonstrate that *E*(*I* |*K*) is an excellent approximation under a single-hits assumption (i.e. w/o replacement) regardless of the underlying network properties (Fig. S1), but tends to overestimate the number of edges that should appear on the DMI network with multiple-hits (Fig. 1); this owes to the evolution of redundant interactions between the same pair of DMI loci (which we call DMI turnover). The edge-accumulation process is nevertheless one corresponding to an accelerating numerical increase, and so the overall snowball effect with respect to edges is robust. While Orr’s (1995) original focus was in deriving the number of hybrid incompatible interactions (edges), it should be noted that the snowball effect applies to loci as well. This observation, though seemingly trivial, necessitates a more detailed accounting of the structure of DMI edges, including how loci are shared, if one wants quantitative results beyond merely edge-counts. That is, given that one has an equation for edge-counts, *E*(*I* |*K*), it does not follow that one has a solid foundation for predicting *E*(*L* |*K*), where *L* are the number of loci implicated in the set of incompatibilities. The set of quantities *I, K, P* lack sufficient information regarding the connections among DMIs necessary for deriving the expectation for DMI-locus counts. Neither do these quantities suffice for predicting the number of DMI-complexes.

More realistic and informative modelling of DMI evolution, therefore, requires substituting Orr’s (1995) uniform probability with a set of combination-specific probabilities (i.e. *P*_ij_, for loci *i* and *j*; and *P*_ij…k_ for higher order combinations) and a mathematical structure capable of expressing information about multilocus connections. The simple pairwise network approach adopted here is a step toward this more fine-grained treatment of the problem, substituting a constant interaction probability with a topology-informed scheme in which only connected loci may interact deleteriously in hybrids (*i.e*., *P* = 0 for unconnected pairs, and fixed *P* otherwise; see Livingstone et al. 2012 for a mathematical treatment aimed at describing the first DMI occurrence).

As divergence progresses in our model, the byproducts of divergence in allopatry are seen to be DMI complexes that build up and aggregate over time, such that the incompatibility network exhibits a massive tangle of interactions in the long run, as opposed to an indefinite build-up of DMI pairs and small complexes. In sum, the snowball effect is accurate for the first several hundred substitutions but, with variation in per-locus evolutionary rates, eventually exhibits a marked deceleration with respect to the numbers of edges and loci contributing to hybrid incompatibility. This deceleration is distinct for each measure (edges, loci), and is influenced most strongly by network density. The snowball effect with respect to DMI complexes (connected components) has a similar approximate accuracy in the first couple hundred generations, prior to exhibiting a negative acceleration (reverse snowball) as distinct complexes fuse together on the incompatibility network. The agglomerative process is opposed mainly by two factors: (1) underlying gene interaction networks generated with the small world algorithm of Watts and Strogatz (1998), and (2) a substitution model featuring the conjunction of (i) multiple hits, (ii) wide variation in substitution rates per locus, and (iii) a strong negative correlation between substitution rates and the numbers of gene interactions per locus.

Our results suggest two factors that can slow down reproductive isolation. First, in low-density (interaction-sparse) networks, the turnover of alleles at already-existing DMIs can result in the turnover of the associated incompatible interactions; the slowdown due to this multiple-hits phenomenon is sizeable for high rates of speciation (see also Olafsson et al. 2016). Second, the size increase of DMI connected components can be interpreted as corresponding to the build-up of DMI genes that affect developmentally correlated hybrid fitness traits; interaction among DMI genes of the same connected component is likely to manifest as diminishing returns epistasis. If the evolution of postzygotic isolation includes saturation effects (i.e. a point where additional substitutions do not sizably reduce hybrid fitness; Welch 2004), perhaps on a modular basis (Orr 1995, Appendix B therein), then the rate of reproductive isolation would be expected to more closely track the accumulation of connected components, rather than the edges, on the incompatibility network. Most notably, the model predicts that the genetic architecture of reproductive isolation in highly diverged species consists largely of a single massive incompatibility cluster (along with a small number of pairs and small DMI complexes).

While our focus has been on randomly generated gene interaction networks of 1000 loci, assuming a biologically realistic network (from Costanzo et al., 2016) recapitulates similar accumulation and agglomeration patterns as were observed from the generated networks (Fig. 5B). Indeed, agglomeration occurred on a similar time scale between the yeast network and the 1%-density networks, despite the 5-fold disparity in node counts and 300-fold difference in edge counts. Interestingly, neither the substitution model (single-vs. multiple hits) nor the presence of substitution rate variation (equal rates vs. per-locus rates proportional to the dN/dS values from Peter et al. 2018) had a strong effect on accumulation and agglomeration outcomes (Figs. 5B, S8). This suggests that DMI network evolution is insensitive to fixation-rate variation among loci, but the true extent of disparity in selective fixation rates across the genome (including expression changes) is not well characterized and may be underestimated by extrapolating from dN/dS alone. (Recall the differential outcomes displayed in Fig. 5A where rate-variation directly reflected vertex-degree variation.)

The pairwise incompatibility network does not exist in isolation from incompatibility networks of higher-order epistasis (representable as hypergraphs). Kalirad and Azevedo (2017) demonstrated that the evolution of complex DMIs systematically influences the trajectory of pairwise DMIs. This is because complex epistasis governs which gene pairs are liable to exhibit incompatibility in the first place. Complex DMIs can thus partially erase the build-up of evolved DMI pairs and may even convert their overall numerical increase to a linear rate. At present, it is unclear to what extent the conflicting influences among levels of epistasis holds genome-wide, outside of the context of RNA-folding (the focus of their article).

As we have shown, the evolution of DMIs is shaped by the interplay between network architecture (topology and density) and substitution dynamics (hit model, rate variation, and rate–degree correlations). This expanded parameter space enables a richer descriptive framework in which DMI loci are not only counted but also tracked by their position and connectivity within a complexifying incompatibility network. An extension to our network approach is possible by considering polymorphism at short-term scales. Many DMIs are not the results of fixed differences but rather are already present in the parent species as variable incompatibilities (Cutter 2011). Because epistasis and recessivity act to mask the expression of deleterious gene combinations, so-called synthetic lethals (and detrimentals, more generally) may segregate at appreciable frequencies and form a basis for the persistence of rare within-species dysfunction (Philips and Johnson 1998; Lachance et al. 2011). Deleterious polymorphism of this sort might lay the foundations for eventual between-species isolation, and a network description of this incipient state of speciation merits elaboration.

---

**Figure S1.**
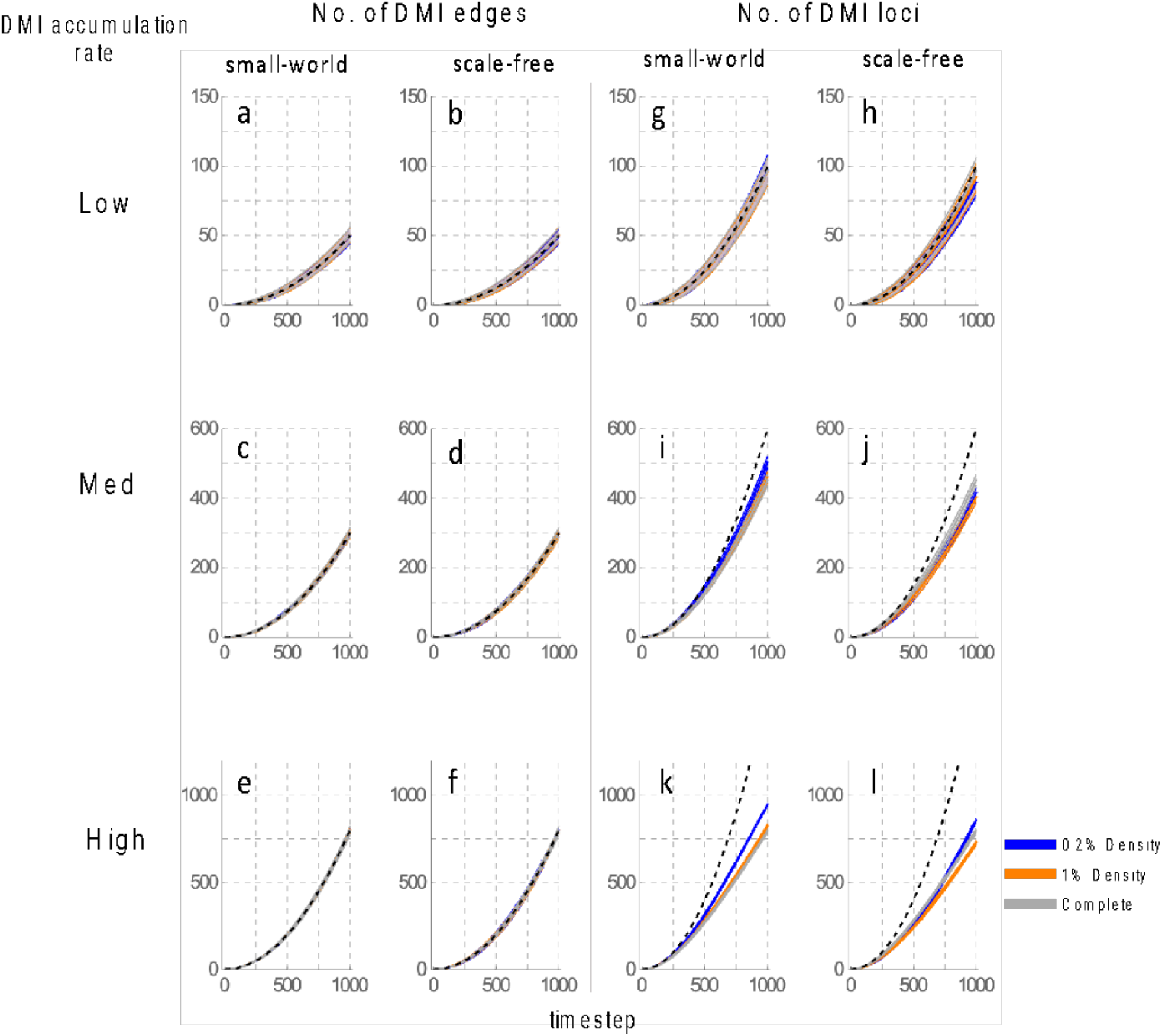
Incompatibility accumulation under a single-hits, equal rates substitution model. Counts of incompatibility edges and loci (means ± 1 mean deviation) are given for the low, medium, and high (density-adjusted) P_inc_ regimes. Accumulation occurred over 1000 timesteps, assuming small-world and scale-free gene interaction networks with 1000 loci. Density of the underlying interaction network is given by color (0.2%– **blue**, 1%-**orange**, Complete – **gray**). Complete graph trajectories are duplicated in corresponding small-world and scale-free panels. (a-f) Dashed line corresponds to Orr’s (1995) binomial expectation for DMI incompatibilities. (g-l) Dashed line gives DMI locus expectation based on a 2:1 ratio of loci to edges.

**Figure S2.**
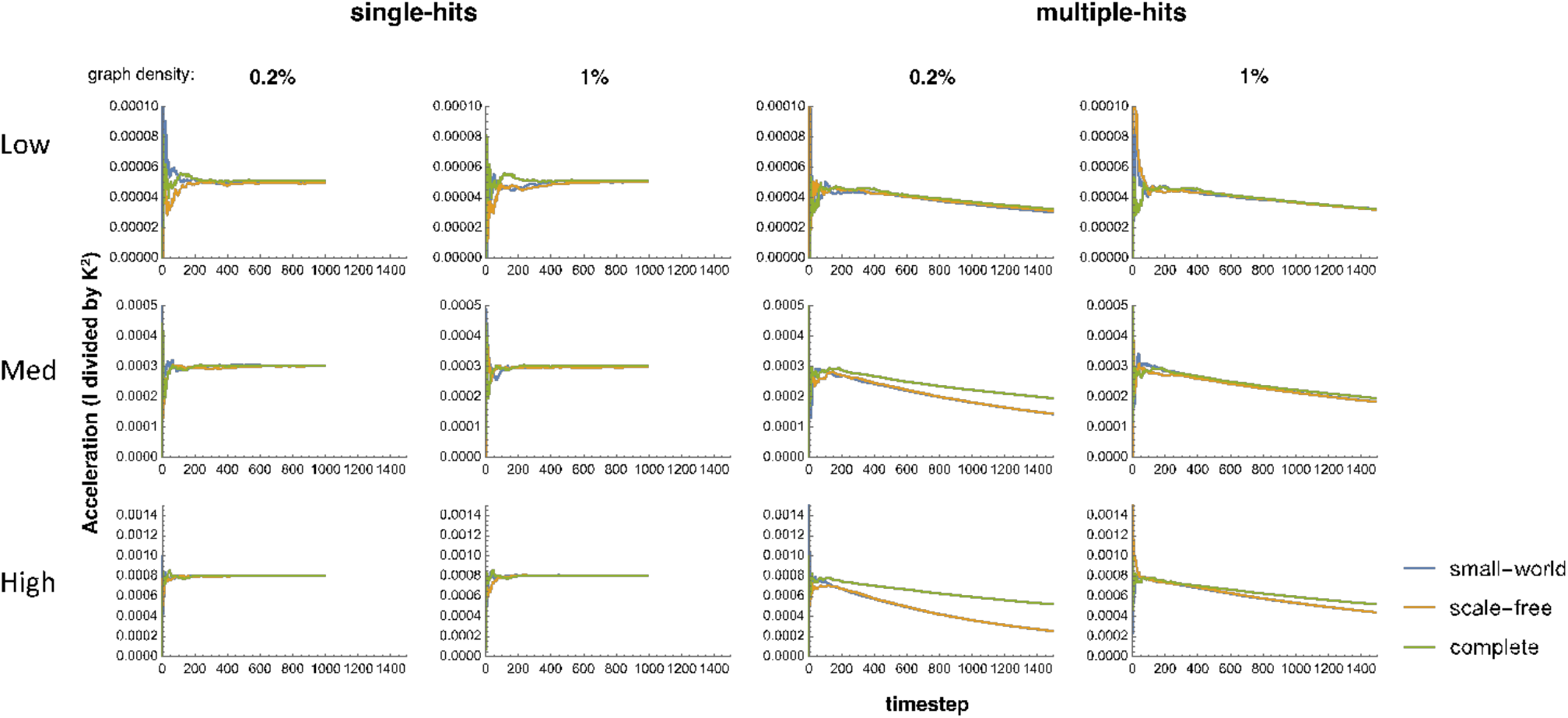
Temporal trend in the DMI accumulation rate. The rate of change between timesteps in the DMI (edge) accumulation rate (referred to as ‘acceleration’ of DMI increase on the vertical axis) is given by I/K (the rate) divided by K, or I/K^2^, where I is the number of edges on the DMI network and K is the number of substitutions. Small-world interaction networks are given in **blue**, scale-free in **orange**, and complete in **green**; blue and orange strongly overlap in the bottom four “multiple-hits” panels. Incompatibility regime is given by the row labels, and graph density by column labels.

**Figure S3.**
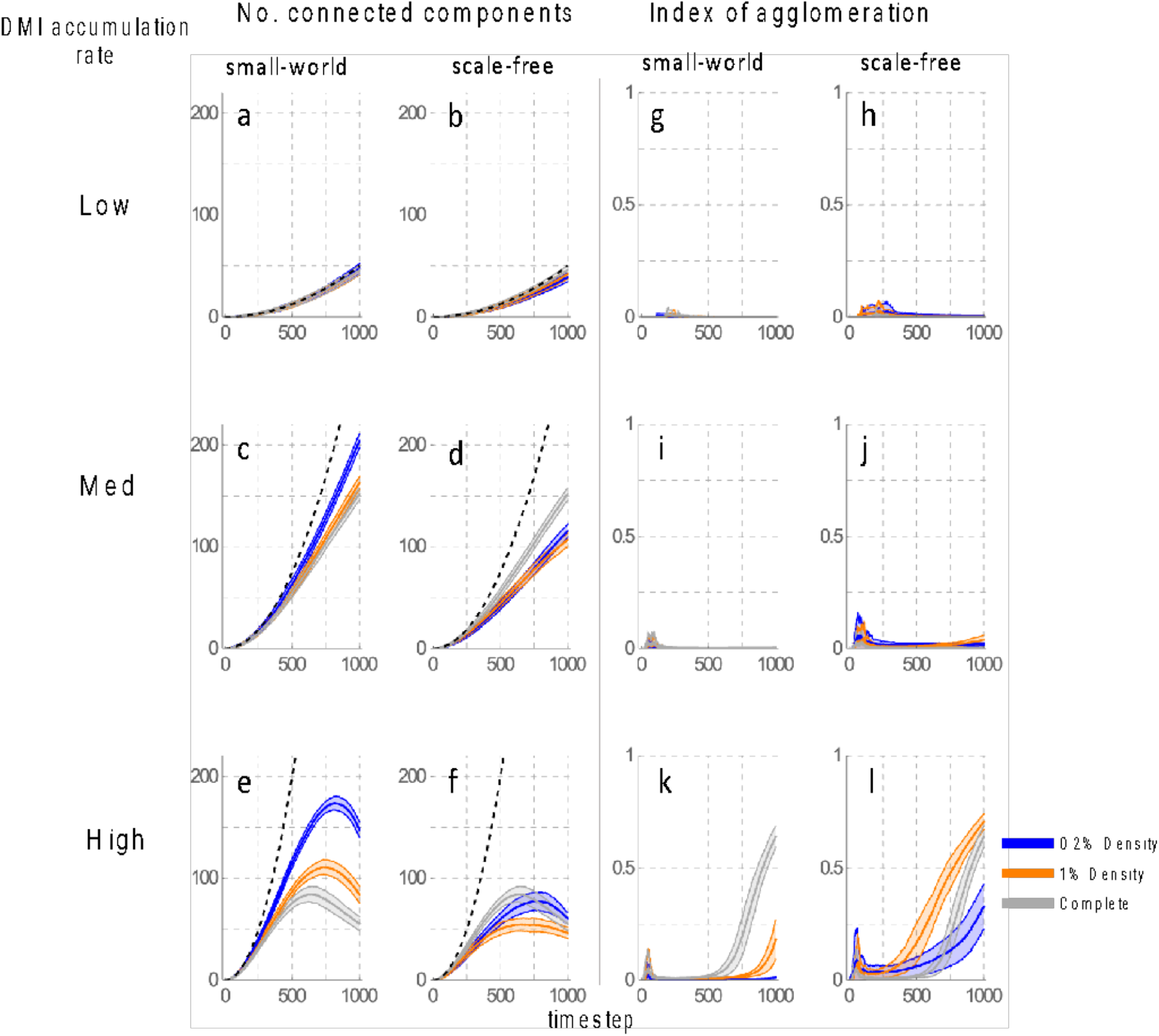
Incompatibility agglomeration under a single-hits, equal rates substitution model. (a-f) The number of connected components on the pairwise incompatibility graph that have 3 or more nodes (means ± 1 mean deviation). Dashed line corresponds to Orr’s (1995) binomial expectiation for DMI incompatibilities. (g-l) I_agg_ (means ± 1 mean deviation). Density of the underlying interaction network is given by color (0.2% edges– **blue**, 1%-**orange**, Complete – **gray**). Complete graph trajectories are duplicated in corresponding SW and BA panels.

**Figure S4.**
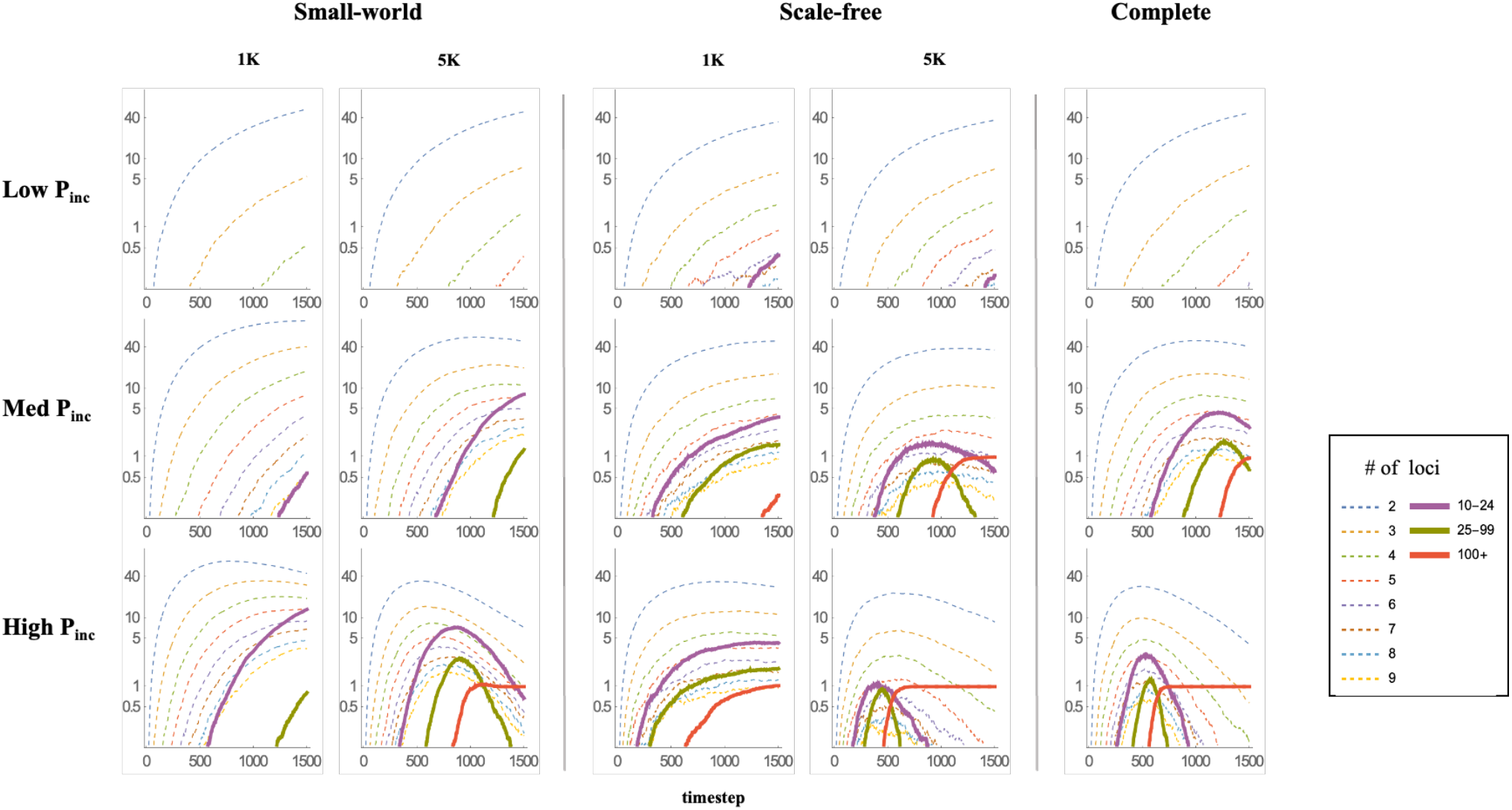
Evolution of the size distribution of agglomeration clusters under a multiple-hits, equal rates substitution model. Mean counts of connected components on the pairwise incompatibility network, broken down by size. Dashed lines correspond to small clusters, while thick, unbroken lines follow the trajectories of larger agglomeration clusters (i.e. connected components with 10 or more nodes). Vertical axis is log-scaled.

**Figure S5.**
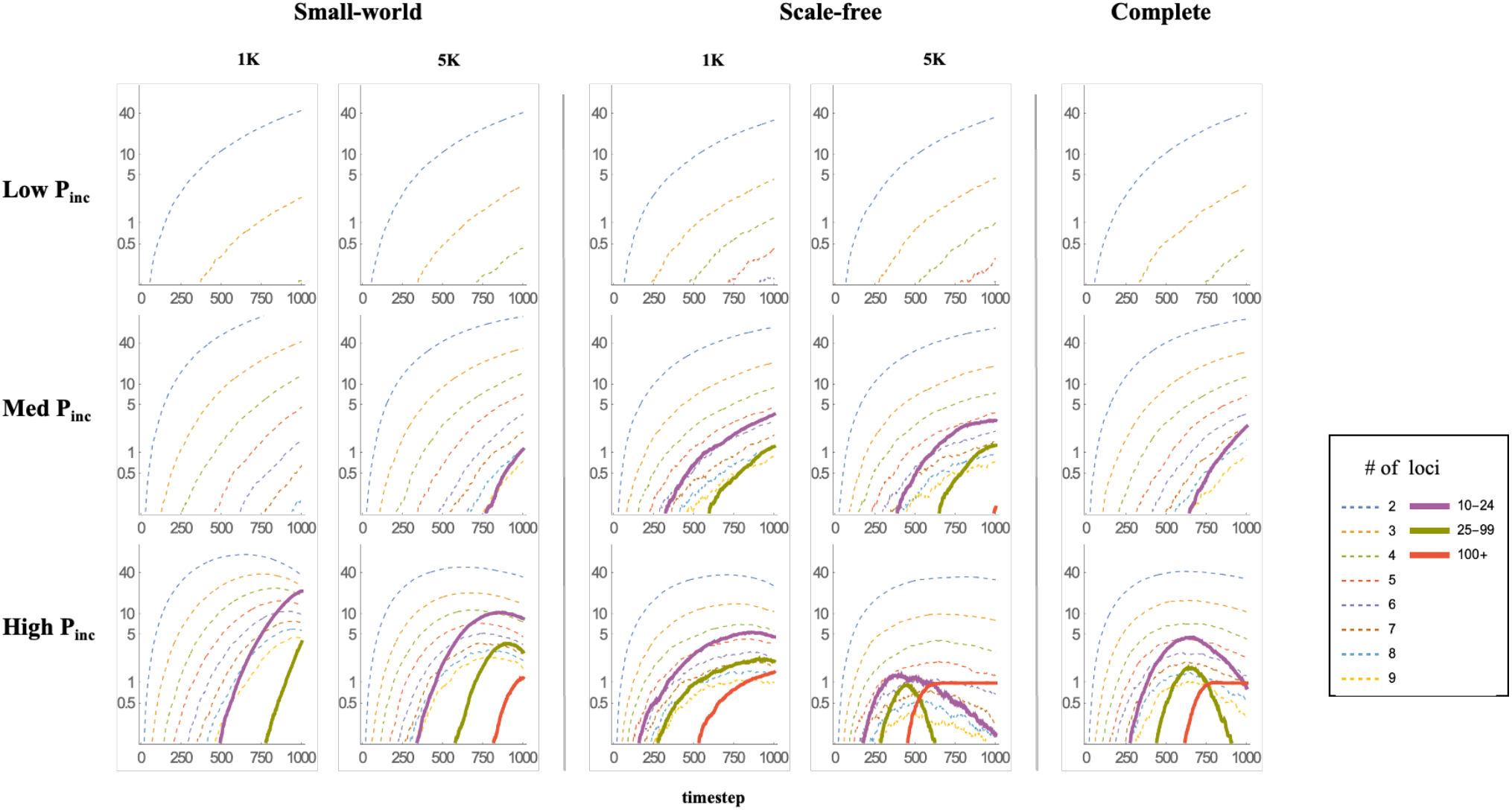
Evolution of the size distribution of agglomeration clusters under a single-hits, equal rates substitution model. Mean counts of connected components on the pairwise incompatibility network, broken down by size. Dashed lines correspond to small clusters, while thick, unbroken lines follow the trajectories of larger agglomeration clusters (connected components with 10 or more nodes). Vertical axis is log-scaled.

**Figure S6.**
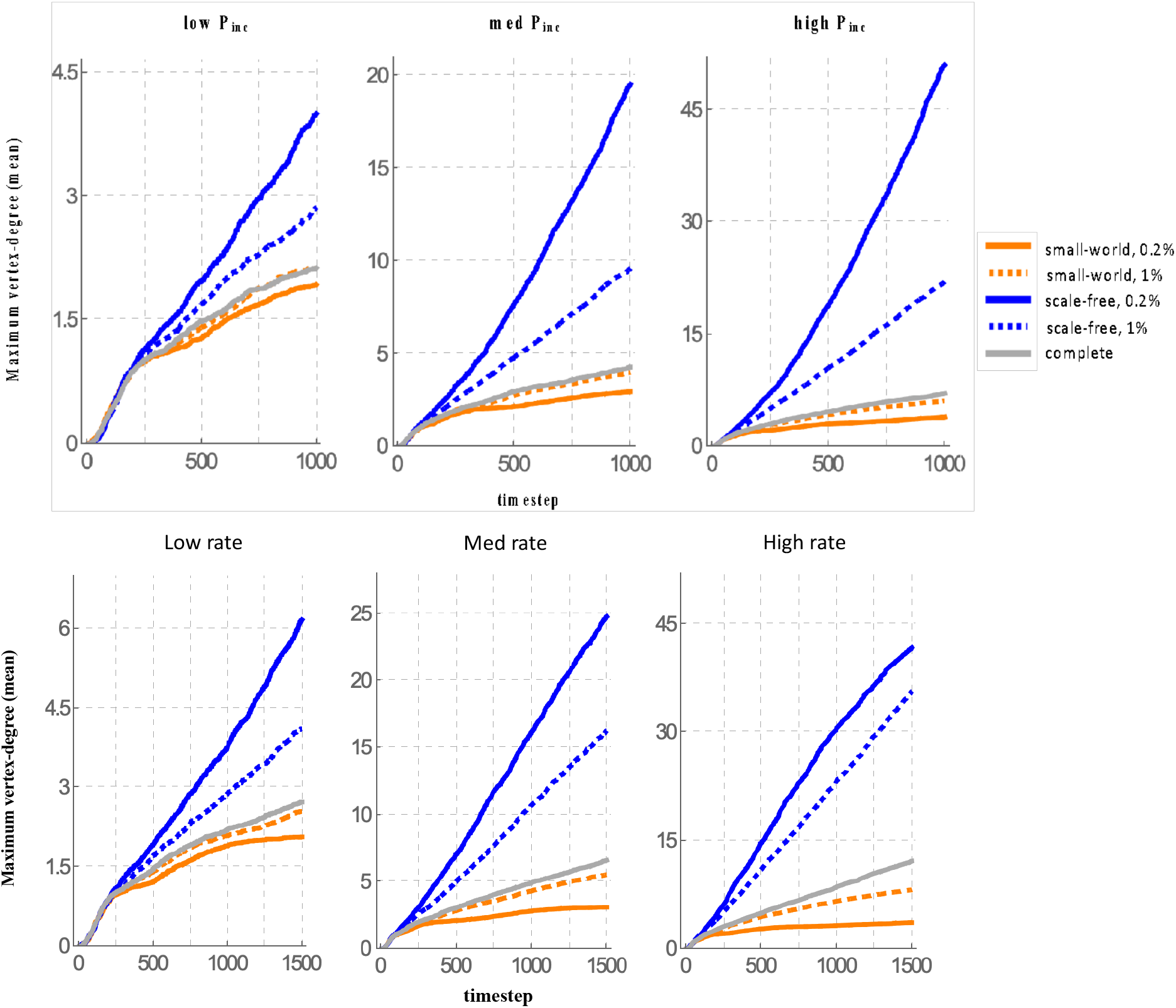
Maximum vertex-degree on the DMI network. The maximum vertex-degree of the incompatibility network is averaged over all 250 replicates for a single-hit model (top row), and a multiple-hits model (bottom row) of substitutions.

**Figure S7.**
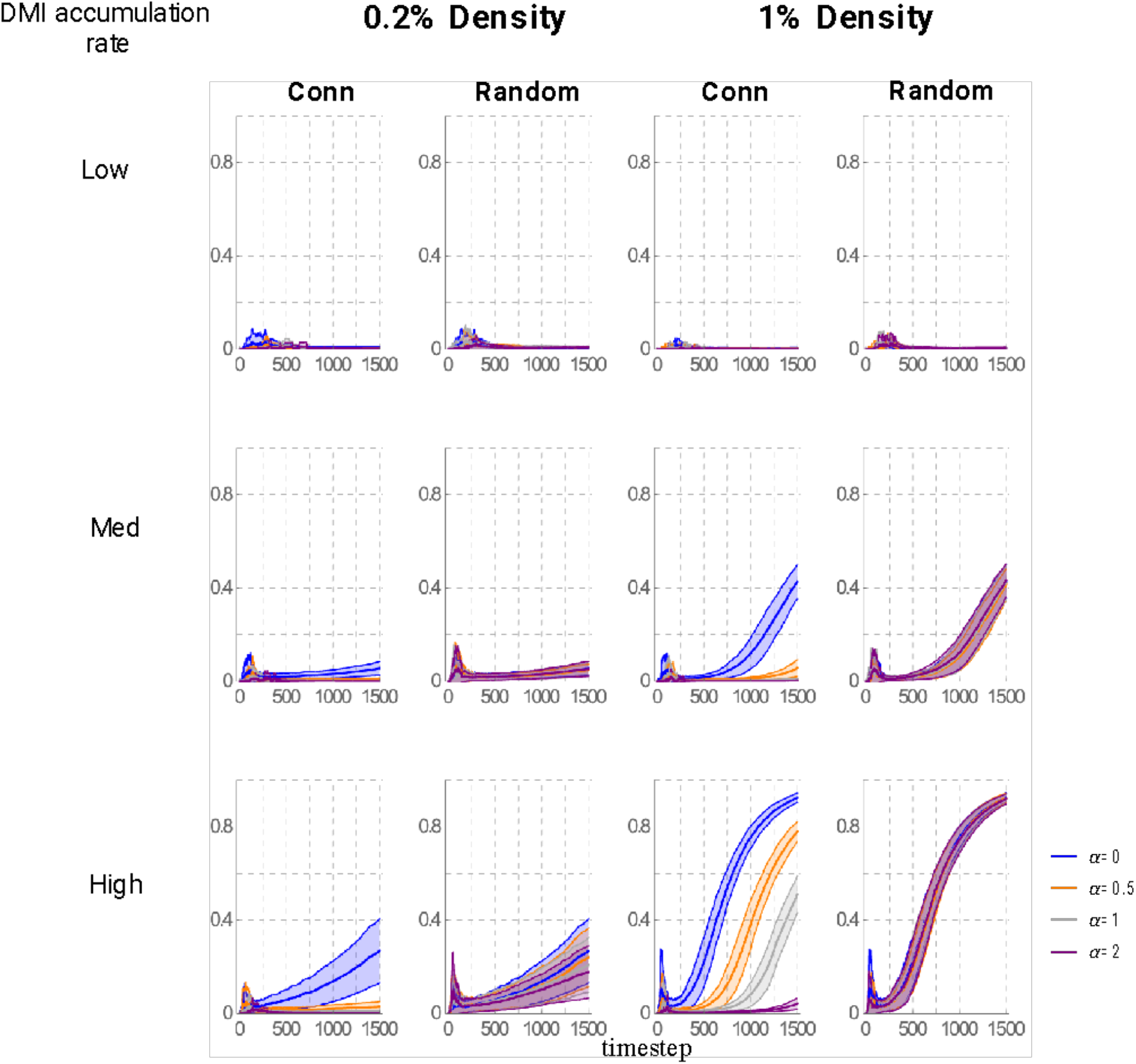
Index of agglomeration under differential substitution rates. Simulation results assuming scale-free networks with 0.2% (left) and 1% (right) density are displayed. Trajectories assume *α* values of 0 (blue; equal rates), 0.5 (orange; weak differences in rates), 1 (grey; moderate differences), and 2 (purple; strong differences). Substitution rates are negatively correlated with degrees under the “Conn” column (connectivity), or are otherwise randomly associated (“Random”).

**Figure S8.**
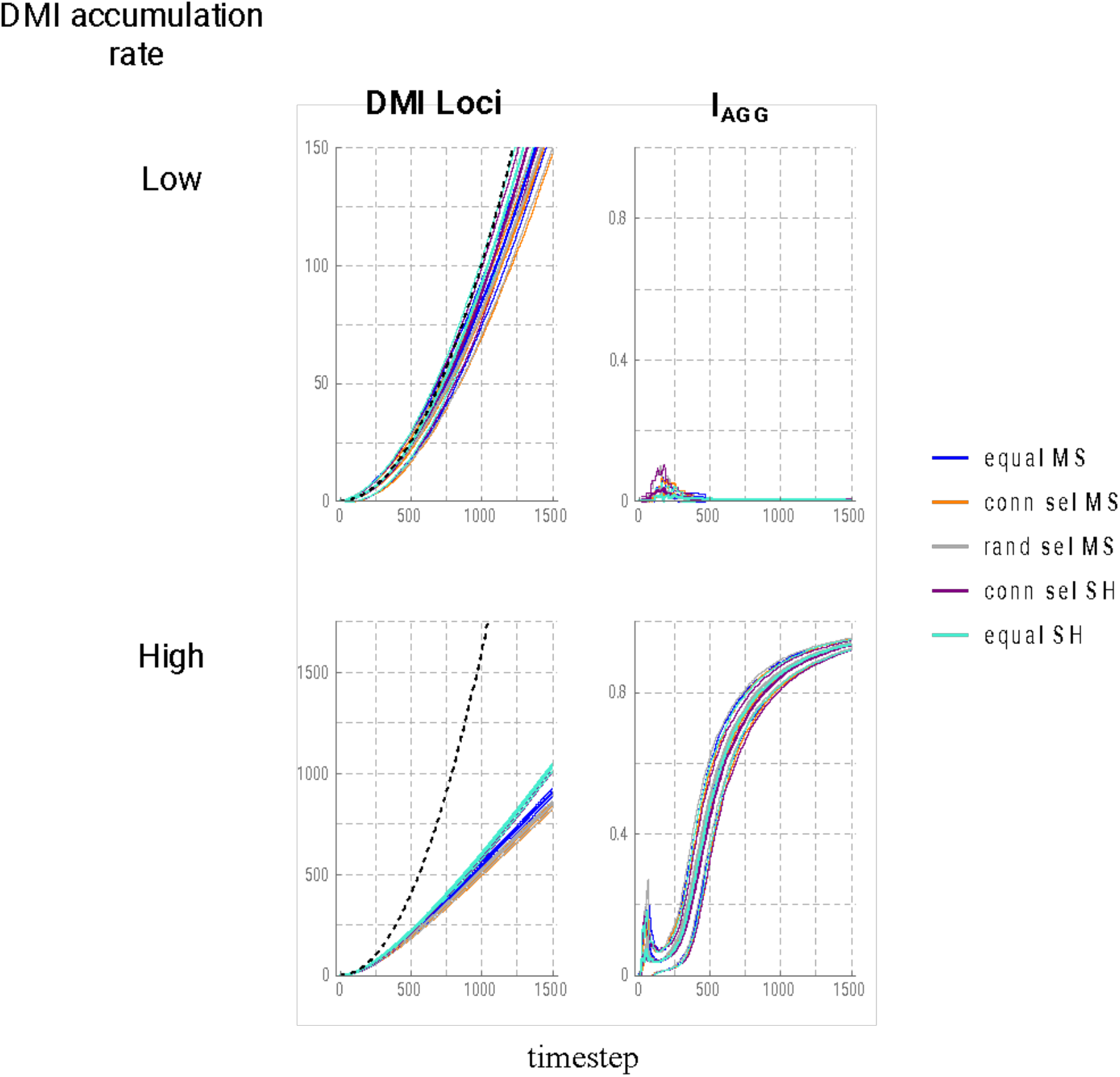
DMI accumulation and agglomeration on the assumption of the yeast gene interaction network. Simulation results assuming the yeast interaction network (see Methods) are displayed for various models of substitution rates. **Equal MS:** equal rates, multiple subs. **Conn sel MS:** substitution rates are proportional to dN/dS values (obtained from Peter et al. 2018), multiple subs. **Rand sel MS:** vector of node labels from Conn sel MS randomized. **Conn sel SH:** substitution rates are proportional to dN/dS values, single-hits. **Equal SH:** equal rates, single-hits. Dashed line for “DMI Loci” is the locus-count expectation based on a 2:1 extrapolation from edges.

## Works Cited

Ayala-López, Julio A., and Claudia Bank. “What can we gain from modelling complex hybrid incompatibilities?” Evolutionary Journal of the Linnean Society 4, no. 1 (2025): kzae034.

Barabási, Albert-László, and Réka Albert. “Emergence of scaling in random networks.” Science 286, no. 5439 (1999): 509–512.

Cabot, Eric L., Andrew W. Davis, Norman A. Johnson, and Chung I. Wu. “Genetics of reproductive isolation in the Drosophila simulans clade: complex epistasis underlying hybrid male sterility.” Genetics 137, no. 1 (1994): 175–189.

Costanzo, Michael, Benjamin VanderSluis, Elizabeth N. Koch, Anastasia Baryshnikova, Carles Pons, Guihong Tan, Wen Wang et al. “A global genetic interaction network maps a wiring diagram of cellular function.” Science 353, no. 6306 (2016): aaf1420.

Cutter, Asher D. “The polymorphic prelude to Bateson–Dobzhansky–Muller incompatibilities.” Trends in ecology & evolution 27, no. 4 (2012): 209–218.

Dobzhansky, T. H. “Studies on hybrid sterility. II. Localization of sterility factors in Drosophila pseudoobscura hybrids.” Genetics 21, no. 2 (1936): 113.

Erdos, Paul, and Alfréd Rényi. “On the evolution of random graphs.” Publications of the Mathematical Institute of the Hungarian Academy of Sciences 5, no. 1 (1960): 17–60.

Heidbreder, Patrick, Noora Poikela, Pierre Nouhaud, Tuomas Puukko, Konrad Lohse, and Jonna Kulmuni. “Genomic incompatibilities are persistent barriers when speciation happens with gene flow in Formica ants.” bioRxiv (2025): 2025–03.

Kalirad, Ata, and Ricardo BR Azevedo. “Spiraling complexity: a test of the snowball effect in a computational model of RNA folding.” Genetics 206, no. 1 (2017): 377–388.

Kao, Katy C., Katja Schwartz, and Gavin Sherlock. “A genome-wide analysis reveals no nuclear Dobzhansky-Muller pairs of determinants of speciation between S. cerevisiae and S. paradoxus, but suggests more complex incompatibilities.” PLoS genetics 6, no. 7 (2010): e1001038.

Koevoets, T., O. Niehuis, L. Van De Zande, and L. W. Beukeboom. “Hybrid incompatibilities in the parasitic wasp genus Nasonia: negative effects of hemizygosity and the identification of transmission ratio distortion loci.” Heredity 108, no. 3 (2012): 302–311.

Kondrashov, Alexey S., Shamil Sunyaev, and Fyodor A. Kondrashov. “Dobzhansky–Muller incompatibilities in protein evolution.” Proceedings of the National Academy of Sciences 99, no. 23 (2002): 14878–14883.

Lachance, Joseph, Norman A. Johnson, and John R. True. “The Population Genetics of X–Autosome Synthetic Lethals and Steriles.” Genetics 189, no. 3 (2011): 1011–1027.

Livingstone, Kevin, Peter Olofsson, Garner Cochran, Andrius Dagilis, Karen MacPherson, and Kerry A. Seitz Jr. “A stochastic model for the development of Bateson–Dobzhansky–Muller incompatibilities that incorporates protein interaction networks.” Mathematical biosciences 238, no. 1 (2012): 49–53.

Matute, Daniel R., Ian A. Butler, David A. Turissini, and Jerry A. Coyne. “A test of the snowball theory for the rate of evolution of hybrid incompatibilities.” Science 329, no. 5998 (2010): 1518–1521.

Moyle, Leonie C., and Takuya Nakazato. “Complex epistasis for Dobzhansky–Muller hybrid incompatibility in Solanum.” Genetics 181, no. 1 (2009): 347–351.

Moran, Benjamin M., Cheyenne Y. Payne, Daniel L. Powell, Erik NK Iverson, Alexandra E. Donny, Shreya M. Banerjee, Quinn K. Langdon et al. “A lethal mitonuclear incompatibility in complex I of natural hybrids.” Nature 626, no. 7997 (2024): 119–127.

Muller, H. “Isolating mechanisms, evolution, and temperature.” Biol. Symp., vol. 6, pp. 71–125. 1942.

Olofsson, Peter, Kevin Livingstone, Joshua Humphreys, and Douglas Steinman. “The probability of speciation on an interaction network with unequal substitution rates.” Mathematical biosciences 278 (2016): 1–4.

Orr, H. Allen. “The population genetics of speciation: the evolution of hybrid incompatibilities.” Genetics 139, no. 4 (1995): 1805–1813.

Orr, H. Allen, and Michael Turelli. “The evolution of postzygotic isolation: accumulating Dobzhansky-Muller incompatibilities.” Evolution 55, no. 6 (2001): 1085–1094.

Palopoli, Michael F., and Chung-I. Wu. “Genetics of hybrid male sterility between Drosophila sibling species: a complex web of epistasis is revealed in interspecific studies.” Genetics 138, no. 2 (1994): 329–341.

Peter, Jackson, Matteo De Chiara, Anne Friedrich, Jia-Xing Yue, David Pflieger, Anders Bergström, Anastasie Sigwalt et al. “Genome evolution across 1,011 Saccharomyces cerevisiae isolates.” Nature 556, no. 7701 (2018): 339–344.

Phadnis, Nitin, EmilyClare P. Baker, Jacob C. Cooper, Kimberly A. Frizzell, Emily Hsieh, Aida Flor A. de la Cruz, Jay Shendure, Jacob O. Kitzman, and Harmit S. Malik. “An essential cell cycle regulation gene causes hybrid inviability in Drosophila.” Science 350, no. 6267 (2015): 1552–1555.

Phillips, Patrick C., and Norman A. Johnson. “The population genetics of synthetic lethals.” Genetics 150, no. 1 (1998): 449–458.

Satokangas, Ina, S. H. Martin, H. Helanterä, J. Saramäki, and J. Kulmuni. “Multi-locus interactions and the build-up of reproductive isolation.” Philosophical Transactions of the Royal Society B 375, no. 1806 (2020): 20190543.

Sherman, Natasha A., Anna Victorine, Richard J. Wang, and Leonie C. Moyle. “Interspecific tests of allelism reveal the evolutionary timing and pattern of accumulation of reproductive isolation mutations.” PLoS Genetics 10, no. 9 (2014): e1004623.

Tang, Shanwu, and Daven C. Presgraves. “Evolution of the Drosophila nuclear pore complex results in multiple hybrid incompatibilities.” Science 323, no. 5915 (2009): 779–782.

Turelli, Michael, and H. Allen Orr. “Dominance, epistasis and the genetics of postzygotic isolation.” Genetics 154, no. 4 (2000): 1663–1679.

Wang, Richard J., Michael A. White, and Bret A. Payseur. “The pace of hybrid incompatibility evolution in house mice.” Genetics 201, no. 1 (2015): 229–242.

Watts, Duncan J., and Steven H. Strogatz. “Collective dynamics of ‘small-world’ networks.” Nature 393, no. 6684 (1998): 440–442.

Welch, John J. “Accumulating Dobzhansky-Muller incompatibilities: reconciling theory and data.” Evolution 58, no. 6 (2004): 1145–1156.

Wolfram Research, Inc., 2021. Mathematica version 13.0. Champaign, IL

